# Phyloformer: towards fast and accurate phylogeny estimation with self-attention networks

**DOI:** 10.1101/2022.06.24.496975

**Authors:** Luca Nesterenko, Bastien Boussau, Laurent Jacob

## Abstract

An important problem in molecular evolution is that of phylogenetic reconstruction, that is, given a set of sequences descending from a common ancestor, the reconstruction of the binary tree describing their evolution from the latter. State-of-the-art methods for the task, namely Maximum likelihood and Bayesian inference, have a high computational cost, which limits their usability on large datasets. Recently researchers have begun investigating deep learning approaches to the problem but so far these attempts have been limited to the reconstruction of quartet tree topologies, addressing phylogenetic reconstruction as a classification problem. We present here a radically different approach with a transformer-based network architecture that, given a multiple sequence alignment, predicts all the pairwise evolutionary distances between the sequences, which in turn allow us to accurately reconstruct the tree topology with standard distance-based algorithms. The architecture and its high degree of parameter sharing allow us to apply the same network to alignments of arbitrary size, both in the number of sequences and in their length. We evaluate our network Phyloformer on two types of simulations and find that its accuracy matches that of a Maximum Likelihood method on datasets that resemble training data, while being significantly faster.

## 1 Introduction

Phylogenies are widely used in biology to convey the history of a group of species or sequences. Most of the time, molecular phylogenies are estimated from aligned nucleotide or amino acid sequences using model-based approaches in the Maximum Likelihood (ML) or Bayesian frameworks, which consider all sequences at once. The models typically are Markovian, and handle events of substitution whereby a nucleotide (respectively amino acid) is replaced by another. Under these models, likelihood computation is made using a costly sum-product algorithm (Felsenstein’s pruning algorithm (Felsenstein, 1981)). Parameters of these models include rates of substitution, the topology of the phylogeny, and its branch lengths. In the Maximum Likelihood framework, inference of continuous parameters is obtained through numerical optimization, and the tree topology is found by exploring the large tree space through local rearrangements. Tree space exploration and parameter optimization involve computing the likelihood numerous times, which make the approach computationally intensive. In the Bayesian framework, inference is typically performed through Markov Chain Monte Carlo methods, in which iterations again require likelihood computations. As a result, in both cases inference is computationally expensive, and a vast amount of work has been devoted to optimizing and parallelizing inference algorithms (Kozlov et al., 2019; Minh et al., 2020; Suchard and Rambaut, 2009; Guindon et al., 2010). Despite these efforts, the footprint of phylogenetic reconstruction can be very large (Jarvis et al., 2014; Philippe et al., 2019; Kumar, 2022), and is expected to grow as the amount of sequence data produced every year keeps increasing.

Faster approaches to phylogenetic reconstruction are often based on pairwise distance methods, which work in two steps. Firstly, a matrix of distances between pairs of sequences is computed using a model of sequence evolution. This distance estimation is typically performed in the maximum likelihood framework, but in this case the likelihood computation is much faster than with the pruning algorithm because only a pair of sequences is analyzed together instead of the whole alignment. Secondly, a clustering algorithm is used to cluster the sequences based on these distances. The Neighbor Joining (NJ) (Saitou and Nei, 1987) algorithm has been widely used for this second step, and is guaranteed to recover the correct topology provided that the input distance matrix is close enough to the real one. Unfortunately, distance approaches are less accurate than ML or Bayesian methods (Guindon and Gascuel, 2003). As a result, they are often confined to the analysis of very large phylogenies, or as starting estimates for more accurate and costly analyses (e.g., Guindon and Gascuel (2003)).

In this context researchers have recently turned to a different inference paradigm based on supervised learning, illustrated, in our case, in Figure 1. This paradigm exploits the fact that simulating data under probabilistic models of sequence evolution is computationally cheap, even in cases where maximizing the likelihood under these models is expensive. Accordingly, large numbers of pairs of phylogenetic trees and multiple sequence alignments (MSAs) evolved along these trees can be generated, and used, via a supervised learning approach, to learn a function which takes an MSA as input and outputs the phylogenetic tree. Learning this function can be a computationally intensive, but once done the function can be used to reconstruct a tree from an MSA very rapidly, regardless of the complexity of the model of sequence evolution that was used to generate the training examples. However, this function has several distinctive features that complicate the supervised learning approach. Firstly, its output is a tree topology and not real values, as output by standard functions used when learning from data. Secondly, its input is an MSA, which can involve any number of aligned sequences of any given length, for which the function must output a tree with the same number of leaves. Thirdly and lastly, the function should provide the same answer regardless of the order of the sequences in the MSA.

**Figure 1:**
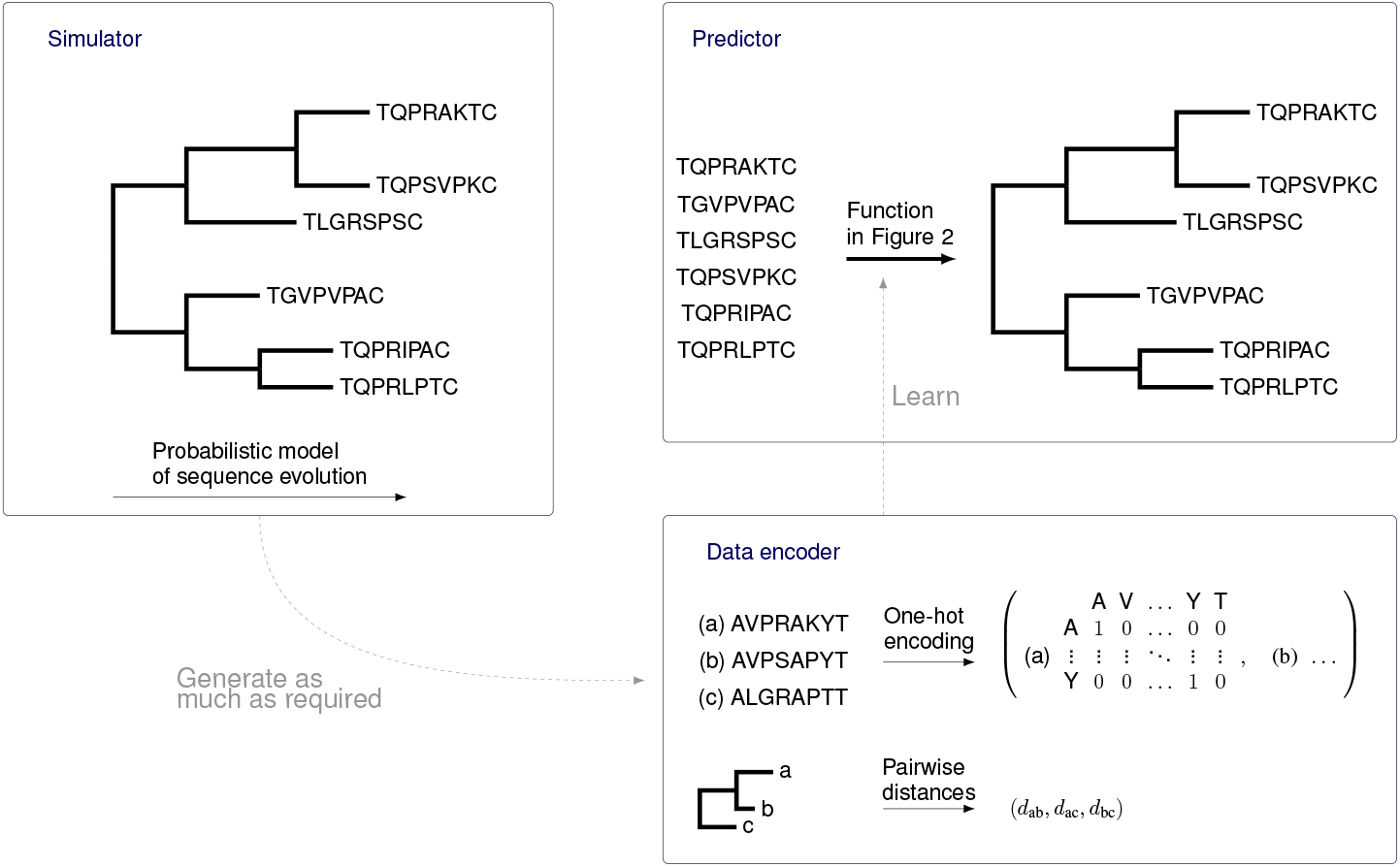
Learning a function that reconstructs a phylogenetic tree from an MSA. We simulate MSAs along known or randomly generated trees (Simulator panel). Once encoded, we use the examples of MSAs and corresponding trees to optimize the prediction function, whose form is detailed in Figure 2

The method that we introduce here works around the first peculiarity by outputting a set of evolutionary distances. If correctly inferred, these distances are enough to recover the tree topology with fast clustering methods such as NJ or BioNJ (Gascuel, 1997). We deal with the two other issues by relying on self-attention, a concept recently popularized in natural language processing (Vaswani et al., 2017), adapting a transformer-based architecture (Rao et al., 2021) for the task. In a nutshell, self-attention builds functions over unordered sets of elements, which would be pairs of aligned sequences in the MSA in our case. The learnable pieces (Figure 2, top right panel) define how each pair representation shares information with the others. This makes the resulting function adaptable to any number of pairs, and equivariant to their permutation—in the sense that permuting the sequences in the input MSA leads to a consistent permutation of the predicted evolutionary distances. The function starts by combining aligned sequences into pair representations, which in themselves are enough to compute *e.g.,* pairwise Hamming distances but that are blind to the other pairs. It then iteratively updates each of them using all pairs in the alignment. The updates are optimized by the supervised learning process to make the outcome as predictive as possible of the true evolutionary distances.

**Figure 2:**
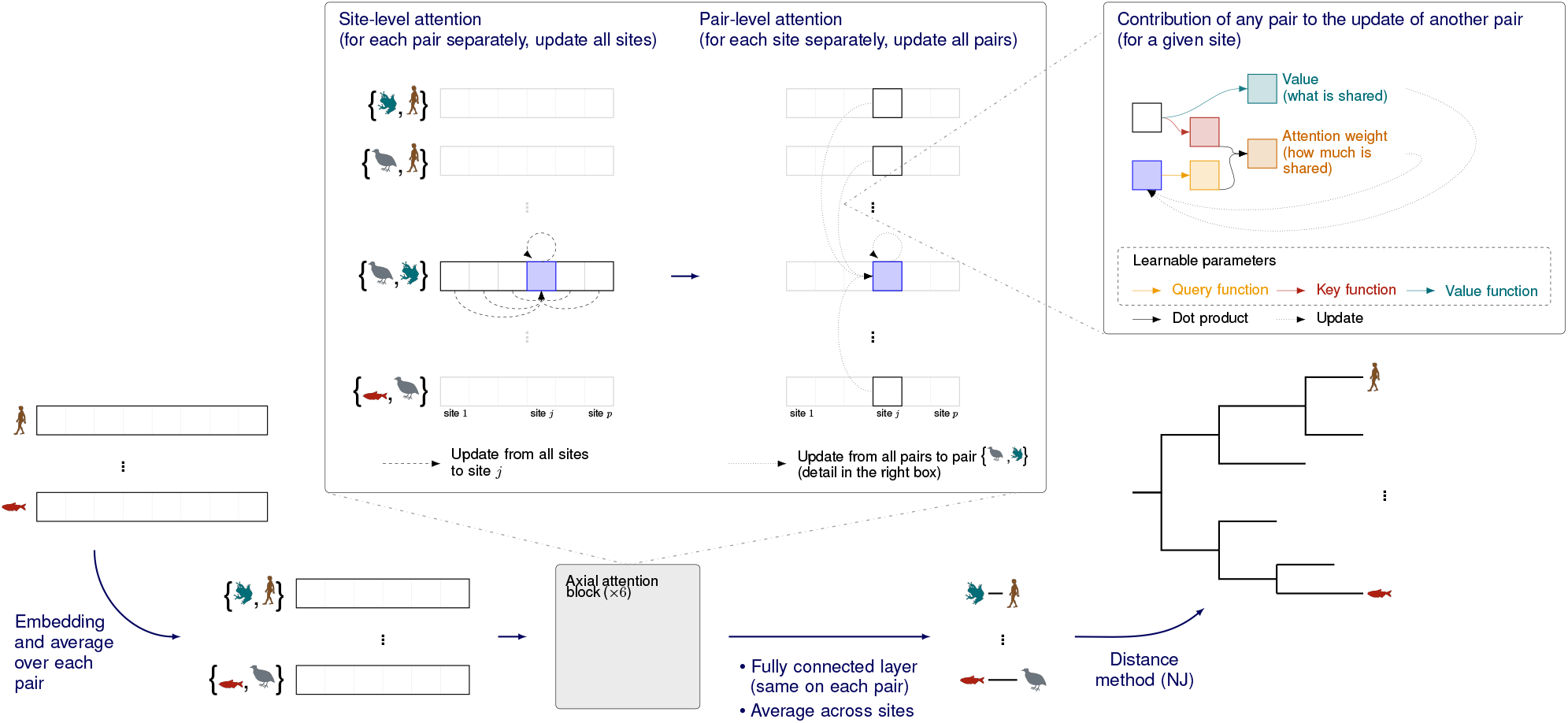
Overview of Phyloformer. Each square denotes a vector of dimension *d* representing one site in one sequence or pair in the MSA, where the value of d can be different at each step. Phyloformer starts (bottom left) from a one-hot encoded MSA, and builds a representation for the pairs (Section 4.1). These pairs then go through several axial attention blocks (Section 4.2) which iteratively build a new representation for each pair that accounts for the entire MSA, by successively sharing information across sites within each pair and across pairs within each site. The sharing mechanism relies on self-attention (right box). We finally use a fully connected feedforward network on each site of the resulting representation and average across sites to predict the evolutionary distance between each pair (bottom right, Section 4.3). These distances can be compared against real distances at training time to optimize the network parameters, or fed to a distance method at test time to reconstruct a phylogeny.

We show on new and previously published simulations that our approach improves on the state of the art of neural networks for phylogenetic reconstruction. Our neural network, named Phyloformer, combined with NJ, consistently outperforms standard distance based methods while being up to 70 times faster than maximum likelihood. Throughout this paper, for ease of notation, we will use the term Phyloformer to refer both to the network itself, that outputs predicted evolutionary distances, and to the corresponding phylogeny reconstruction method that uses these to reconstruct a tree with NJ, the case being clear from the context. The performances of the network combined with different distance-based clustering algorithms, such as BioNJ will be explored in future work.

## 2 Related work

In several fields of bioinformatics, neural networks have been used to extract information from sequence data. Beyond methods that analyze one sequence at a time (Zhou and Troyanskaya, 2015; Alipanahi et al., 2015; Zaheer et al., 2020; Ji et al., 2021), we focus on methods that handle a group of related sequences together, in population genetics, phylogenetic inference and protein structure prediction.

### 2.1 Population genetics

Over the past five years, several neural network methods have been introduced in population genetics. These methods use multiple sequence alignments as input, but do not estimate the phylogeny relating the sequences. Instead, they predict parameters such as effective population size (Sanchez et al., 2021), position and characterize selective sweeps (Flagel et al., 2018) or recombination hotspots (Chan et al., 2018; Adrion et al., 2020). Each of these problems amounts to estimating a global value across the aligned sequences, which does not depend on their order. Consequently, population genetics methods have proposed prediction functions that were invariant by sequence permutation, i.e. whose output does not depend on the order of the sequences in the input alignment (section 4.6).

Of note, the approach used by Sanchez et al. (2021) to achieve invariance bears some similarity to self-attention. Instead of concatenating a global average, self-attention replaces each sequence by a weighted average of some embedding of all sequences in the alignment. Both the embeddings and the weights are optimized during the learning process to make the weighted average as informative as possible to the prediction task. In this sense, attention allows for more expressivity than a uniform average, by learning which sequences should contribute most to the update of which other sequences.

### 2.2 Phylogenetic inference

Few methods have been proposed to learn functions reconstructing phylogenies from simulated alignments. To our knowledge, all of them formulate the problem as a classification across possible topologies. Therefore, given the super-exponential growth in the number of sequences of the number of possible unrooted tree topologies, these methods are restricted to trees with four leaves (quartet trees), that then need to be combined to obtain larger trees. This combination of quartet trees can be achieved using methods developed in the late 1990s and early 2000s (e.g., Strimmer and Von Haeseler (1996)), whose accuracy was found to be lower than that of ML or distance methods (Ranwez and Gascuel, 2001). Two methods were the first to reconstruct quartet trees based on biological sequences, (Suvorov et al., 2019), which used a convolutional network similar to that used in Flagel et al. (2018), and Zou et al. (2020), which applied a similar scheme for protein sequences. In their work a convolutional network architecture takes as input an alignment of four sequences having fixed maximum length, and outputs the computed probabilities for each of the three possible unrooted quartet tree topologies. Zaharias et al. (2022) showed that the accuracy of this network was lower than that of ML or distance methods when evaluated on difficult problems involving long branch lengths and shorter sequences (200 sites), for both quartet trees and trees with 20 leaves. More recently, while still considering only quartet trees, Solis-Lemus et al. (2022) proposed a network that requires fewer training samples (Section 4.6), and reported accuracies similar to Zou et al. (2020).

### 2.3 Protein structure prediction

Finally, our work is also methodologically connected to the recent corpus of methods predicting contact between pairs of residues from MSAs, a crucial step in protein structure prediction. Indeed usually the goal of these methods is to predict distances between sites (columns in the alignment) which can be seen as a transposed version of the problem in our framework, that is predicting a distance between sequences (rows in the alignment). This corpus includes the Evoformer module of the most recent version of AlphaFold (Jumper et al., 2021), which also works with a pair (of sites) representation, along with one of the full MSA, employing an attention mechanism, and the MSA transformer (Rao et al., 2021). Our method uses axial attention blocks (Ho et al., 2019) similar to those employed in the latter. However in our case these blocks act on pairs instead of individual sequences and predict an outcome directly from these pair representations. Our network is trained end-to-end to make this prediction, whereas the one by Rao et al. (2021) is pre-trained on a masked language modeling task to learn a data representation that is then used as input for residue contact prediction learning.

## 3 Background: models of sequence evolution and distance based methods

Sequence evolution is classically modeled as a continuous-time Markov chain (Nielsen, 2006; Felsenstein, 2004), whereby substitution probabilities only depend on the current state and not on previous ones. Furthermore, a widespread assumption is that the process is independent and identically distributed (i.i.d.) across sites of the sequence. Therefore, the evolution of a protein sequence of length *L* is modeled by *L* i.i.d. Markov processes running along the branches of a common underlying phylogenetic tree, with the twenty amino acids alphabet as state space 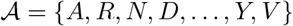. Parameters include the phylogenetic tree, which is assumed without loss of generality to be binary, positive branch lengths, which can be seen as the expected numbers of substitutions per site, exchangeabilities, which characterize the flow of the Markov processes between states, and amino acid profiles, namely the stationary distributions of the Markov processes. More complex models allow e.g., site-wise rates of evolution, which act as multipliers for all branch lengths (Yang, 1994), site-wise or branch-wise changes in equilibrium frequencies (Lartillot and Philippe, 2004; Blanquart and Lartillot, 2006), or non-independent evolution across sites (Robinson et al., 2003).

Probabilistic models can be used to simulate sequence evolution, to reconstruct phylogenies in the Maximum Likelihood framework thanks to topological search and numerical optimization algorithms, or to estimate an evolutionary distance between two sequences. Such distances can then be used for reconstructing a phylogenetic tree as explained below.

A binary tree with positive branch lengths naturally defines a distance matrix on the set of its leaves as

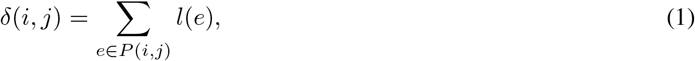

with *l*(*e*) being the length of an edge and *P*(*i,j*) the set of edges on the path from leaf *i* to leaf *j* along the tree. For the converse problem of converting a distance matrix into a tree, the neighbor joining algorithm (Saitou and Nei, 1987) has been a very popular approach. It is guaranteed, given these pairwise distances, to reconstruct the original tree topology. Moreover neighbor joining has *l*_∞_ radius 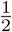 (Atteson, 1999), meaning that the correctness of the reconstructed topology from pairwise distance estimates is mathematically guaranteed as long as these are at most half the shortest edge length in the tree away from their true value. Nevertheless it has been observed that often the algorithm is successful even when this sufficient condition is not met (Mihaescu et al., 2009). To infer all pairwise evolutionary distances, one can use maximum likelihood estimates according to a specific model of sequence evolution. Another approach is to simply take the fraction of differing sites between sequences (the normalized Hamming distance). However this fails to take into account the possibility of multiple substitutions occurring at the same site and leads to underestimating the correct evolutionary distances for long branches or fast evolving sites. Of note, this drawback is shared with another approach that does not explicitly rely on a probabilistic model of sequence evolution, maximum parsimony (Farris, 1970; Fitch, 1971). Overall, distance methods are fast, but consider only two sequences at a time instead of the whole alignment. This probably explains in part why distance methods are empirically less accurate than full scale ML approaches.

## 4 Methods

In our work we draw inspiration from distance methods, but benefit from the self-attention mechanism to use the information from all sequences in the input alignment. Our supervised learning approach therefore goes as follows: for each training sample we simulate evolution along a tree with a given sequence simulator, producing an MSA which, once one-hot encoded, is given in input to the network. The corresponding label is a vector containing all the pairwise distances in lexicographic order, the loss optimized during training being the mean squared error (MSE) between the input label and the output predicted distances. We thus have, for an MSA of *n* sequences of length *L*, an input of size 22 × *n* × *L* and an output of size 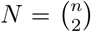. We rely on the paradigm represented in Figure 1: we simulate a large number of MSAs from known or randomly generated trees with probabilistic models of sequence evolution, and use this data to learn a function that takes as input an MSA and attempts to predict the tree along which it evolved. For the purpose of this work, we assume that such a probabilistic model of sequence evolution (left box Simulator) is available to us and focus on how the prediction function (right box Predictor) can be parameterized and how the parameters can be optimized to yield the best possible phylogeny reconstructions. Our approach thereon is summarized in Figure 2, and we now provide detail on each step, starting from a set of MSAs and corresponding phylogenies. Throughout our network, each MSA is successively represented as a *d* × *N* × *L* tensor with each slice in ℝ^d^ representing one position in one of the sequence pairs. The network initially encodes this position and pair independently of all others (vector *φ*^(0)^ in section 4.1), and gradually incorporates information about the rest of the MSA, both across sites and sequences (section 4.2).

### 4.1 From MSA to independent pair representations

We start from a one-hot encoding of aligned sequences (Figure 1, bottom box Data encoder): every sequence *x* is represented as a matrix *φ*^(0)^ (*x*) ∈ {0,1}^22×*L*^ in which each column contains a single non-zero element 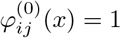, whose coordinate *i* ∈ {1,…,22} denotes the amino acid or gap present in sequence *x* at position *j*. We then represent each pair (*x, x*′) of sequences in the same MSA with the average of their individual representations and write with a slight abuse of notation 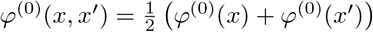. Because *φ*^(0)^(*x,x*′) = *φ*^(0)^(*x*′, *x*), we ensure that the representation is invariant by permutation of the two sequences in the pair.

### 4.2 Axial attention builds MSA-aware pair representations

Self-attention has been recently popularized in the context of natural language processing (Vaswani et al., 2017) and since then has been proven to be among the most successful methods for learning from sequential data (Zaheer et al., 2020; Avsec et al., 2021) both in the context of natural language (Devlin et al., 2018; Brown et al., 2020) and biological sequences (Rives et al., 2021; Elnaggar et al., 2020).

Concretely, self-attention defines a mechanism to update one object *z_i_* using all elements in a set {*z_j_*}_*j*=1,…,*M*_ through a (*query, key, value*) triplet of functions acting on individual objects (schematized in Figure 2, upper right box). The update replaces *z_i_* by a weighted average of all *value*(*z_j_*), where the weights are computed from *query*(*z_i_*) and the corresponding *key*(*z_j_*). The *value* function therefore defines what information on an object *z_j_* should be provided to others, and the interaction between *query* and *key* determines how much is shared from a given *z_j_* to another *z_i_*. For example, the popular dot-product attention used in Vaswani et al. (2017) relies on linear embeddings for vectors *z* ∈ ℝ^*d*^ to define a query *q*, a key *k* and a value *v*:

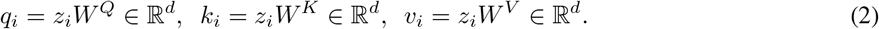

Denoting *K* ∈ ℝ^*d*×*M*^ the matrix whose columns are the (*k_j_*)_*j*=1,…,*M*_, it updates every element *z_i_* by 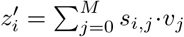, where 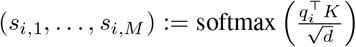 simply contains dot products between *q_i_* and every *k_j_*, scaled by 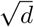 and forced to sum to one by the softmax function. Several such updates can be recursively applied to the *M* elements, progressively sharing information across them. Finally, often what is used in practice is multi-head self attention, which conceptually allows to focus on several different features of the inputs. It is realized using *h* so-called attention heads, that is *h* different triplets 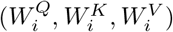, *i* ∈ {1… *h*}, each computing a set of outputs with the aforementioned attention mechanism. The outputs corresponding to each input and the different attention heads are then concatenated and multiplied by a common *matrix W^O^*.

Because the *query, key* and *value* functions act on single objects, they define a flexible recipe to share information across elements of a set, regardless of their order or number. In our context, using dot-product attention on sequence pairs (*z_j_*)_*j*=1,…,*M*_ within MSAs therefore allows learning linear embeddings across alignments with potentially different numbers of sequences, to build new MSA-aware pair representations. The learning process optimizes the weights 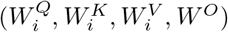 of the embeddings to make, in our context, the final representation as predictive as possible of the corresponding evolutionary distance (see Section 4.3).

Nonetheless, linear embeddings taking in input entire one-hot encoded pairs *φ*^(0)^(*x*, *x*′) ∈ ℝ^22×*L*^ are not ideal because they would learn specific weights for each site in the alignment. Following Rao et al. (2021), we rely on the axial attention technique proposed by Ho et al. (2019); Child et al. (2019) (Figure 2, central box). Every axial attention block takes as input a representation *φ*^(*l*)^(*x, x*′) of every pair in the MSA. It then alternates between updating the representation of each pair separately by sharing information across sites, and updating the representation of each site separately by sharing information across pairs, to output a new representation *φ*^(*l*+1)^(*x, x*′) of the same set of pairs. Both site- and pair-level updates use the same self-attention mechanism, which, acting in the same way on each position-sequence pair representation, regardless of their order or cardinal, allows us to apply the method to MSAs with an arbitrary number of sequences of arbitrary length.

### 4.3 From pair representations to phylogenies

The *r* axial attention blocks of Phyloformer output for every pair of the MSA a tensor *φ*^(*r*)^ (*x, x*′) ∈ ℝ^*d*×*L*^ informed by all pairs in the same MSA. In the end we convert this representation into a single estimate of the evolutionary distance between *x* and *x*′ with a position-wise fully connected layer (section 4.5) with one output feature followed by an average over the sites. Phyloformer is trained end-to-end: the evolutionary distance estimates obtained on simulated MSAs are compared against the true evolutionary distances obtained from the corresponding phylogeny (Figure 1, Data encoder box) using a mean squared error loss function, and the errors are backpropagated through the network to compute a gradient of this loss with respect to all trainable parameters. We use this gradient to improve the parameters, including the attention embeddings, and iteratively make the output distances as close as possible to the true distances. At test time, the output distances are fed to a neighbor joining algorithm (Saitou and Nei, 1987) to reconstruct a phylogeny.

### 4.4 Scalable Phyloformer

A known limitation of self-attention is its poor scalability: the dot-product approach described in Section 4.2 for example requires to compute and store attention weights sij for every pair of elements—each element being either a site or a pair of sequences in the MSA in our context. As a consequence, the axial approach of 4.2 has complexity 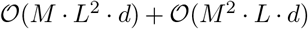 both in memory usage and number of operations. In particular, computing the gradients with respect to the embeddings (*W^Q^, W^K^, W^V^*) by backpropagation requires to store a quadratic number of attention weights *s_ij_* which in our experience led to a bottleneck in training speed even for moderately long sequences *L* = 200 and number *N* = 20, limiting the maximal batch size that could be used during training.

To make the model more scalable we resort to a linear version of self-attention, both for site- and pair-level attention.

#### 4.4.1 Linear self-attention

Following the important success of self-attention, many solutions have been proposed to make this complexity linear in the number of elements in the set (Katharopoulos et al., 2020; Kitaev et al., 2020; Wang et al., 2020; Zaheer et al., 2020). Here we discuss the pair-level self-attention which requires combining *M* pairs, but the same reasoning applies to site-level attention. We rely on the approach of Katharopoulos et al. (2020), who notice that the softmax weights 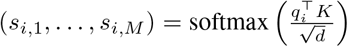 can also be written with inner products 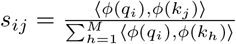 for some non-linear infinite-dimensional mapping 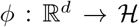 to a Hilbert space 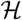 (Schölkopf and Smola, 2002). They propose to replace *ϕ* by some other non-linear, finite-dimensional mappings 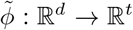, which allows re-writing the update 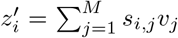 as 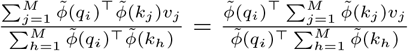. Because each of the two sums can be pre-computed and re-used for every query, this simple factorization reduces both the number of operations and memory usage from 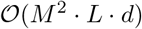 to 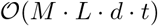.

We follow Katharopoulos et al. (2020) and use an ELU-based mapping (Clevert et al., 2016)

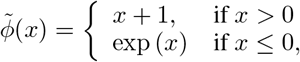

where the operation is applied entrywise, yielding 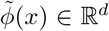 vectors for *x* ∈ ℝ^*d*^. In our experiments, we typically use *d* = 64, making the linear self-attention approach more efficient for pairs (resp. sites) for MSAs with more than *N* =12 sequences (resp. *L* = 64 sites). Consistently with the results reported in Katharopoulos et al. (2020), the linear self-attention strategy provided comparable prediction performances as the regular softmax approach, but sped up the training of Phyloformer by an order of magnitude.

### 4.5 Network architecture and training

The axial attention blocks are based on the MSA transformer which in turn closely follows the original transformer encoder architecture, the main difference here being the use of the much more efficient linear self-attention. All models used in these experiments used *r* = 6 axial attention blocks. Each block used multi-head self-attention (Vaswani et al., 2017) with *h* = 4 heads, successively over the rows and columns of the MSA, followed by a position-wise feed-forward network which acts separately and identically on each position-sequence pair representation, this can also be seen as two consecutive 2D convolutions with 1 × 1 kernels^2^. Each of these three layers were preceded by Layer normalization (Ba et al., 2016) and completed by a skip connection (He et al., 2016). A dimension of *d* = 64 for each position in each pair is used both in input and output of the attention blocks, with the weight matrices 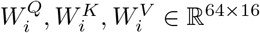 and *W^O^* ∈ ℝ^64×64^. The final position-wise feed-forward network inside the attention block has two layers with a hidden dimension of 256 and a GELU (Hendrycks and Gimpel, 2016) activation function. Next to last, before the average over sites, we have a final position-wise fully connected layer with a softplus activation (Zheng et al., 2015) to constrain the outputs to be positive. The whole architecture is represented in Figure 3. The models were trained with the Adam optimizer (Kingma and Ba, 2015), a learning rate of 4 × 10^-4^ and a batch size of 12, on a single 16GB Nvidia Tesla v100 GPU for 40 hours. To reduce the computational burden we used automatic mixed precision representation of floating points as implemented in PyTorch (Paszke et al., 2019).

**Figure 3:**
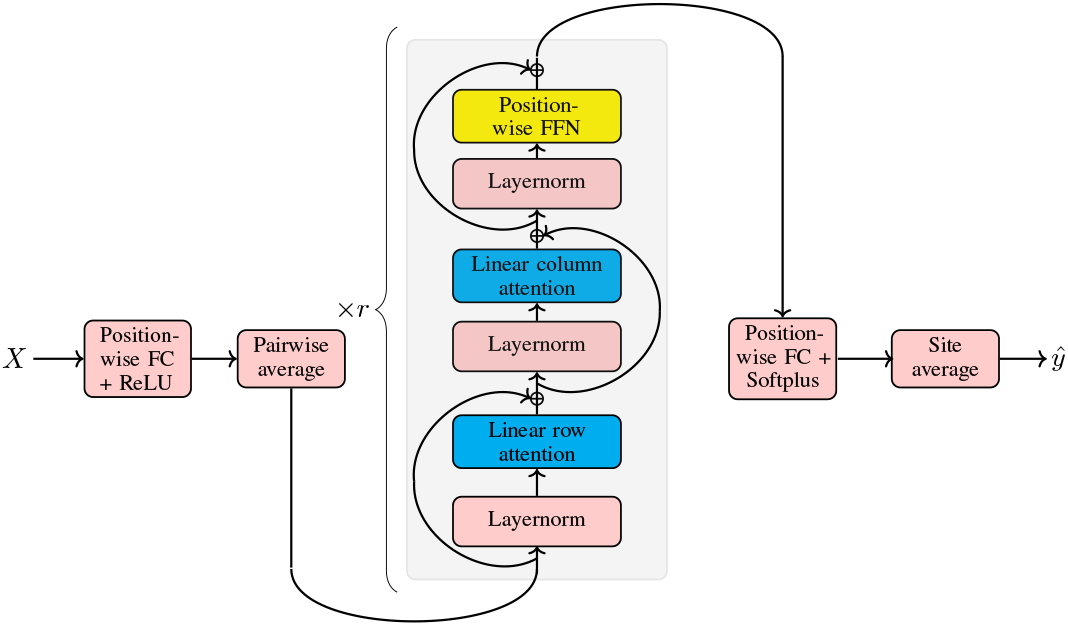
Network architecture of the Phyloformer model. The one-hot encoded amino acids (a vector in ℝ^22^ for each position-sequence pair in the input alignment) are embedded in ℝ^*d*^ via a position-wise fully connected layer, taking pairwise averages over sequences we get the representation in ℝ^*d*×*N*×*L*^ of all the pairs of sequences in the alignment. Up to here each encoded amino acid is processed independently, the subsequent axial attention blocks then allow them to interact building a context aware representation. These multiple blocks are followed by a last position-wise fully connected layer with a softplus activation and a single output feature. Finally averaging over sites gives us the final network prediction.

### 4.6 Invariance and equivariance

It is often desirable that specific transformations of the input data have no effect on the value returned by a network. To this end, one can exploit *symmetries* of the problems at hand *(cf.* Geometric Deep Learning (Bronstein et al., 2021)). We exploit two such symmetries for phylogenetic reconstruction. Firstly, the order of the input sequences is arbitrary, therefore the same evolutionary relationships should be inferred irrespective of which permutation of the sequences is used. Secondly, we train our model using data generated under a site-independent model: as a consequence the same evolutionary relationships should be inferred irrespective of which permutation of the sites is used.

Several expedients have been devised to deal with permutations of sequences inside an MSA. To this end, within the context of population genetics, Flagel et al. (2018) sorted aligned sequences by similarity before feeding them to a non invariant function, while Chan et al. (2018); Sanchez et al. (2021) directly devised permutation-equivariant prediction functions. More specifically, Chan et al. (2018) used a so-called exchangeable network that first applied the same function—a one-dimensional convolutional neural network—to each sequence and then applied a symmetric function—such as the average across sequences—at each site. Sanchez et al. (2021) chose a more progressive approach where they also applied the same convolutional network to each sequence, but iteratively concatenated to the obtained representation of each sequence the average over the current representation of all sequences in the alignment. This process makes all sequences converge to a common representation across iterations.

In the context of phylogenetics, while Zou et al. (2020) resort to data augmentation by considering all the permutations of the input sequences both during training and at inference time, as pointed out in Solis-Lemus et al. (2022), which proposed a network whose output is unchanged when the former is faced with a permutation of the clades for which the unrooted quartet tree topology is invariant, explicitly exploiting the symmetries of the problem instead allows using fewer parameters and training examples.

Phyloformer predicts the set of evolutionary distances between all pairs of sequences. By contrast to the population genetics context, and to phylogenetic reconstruction cast as a classification problem, our prediction function therefore needs to be equivariant instead of invariant: if it returns values *d_ab_, d_ac_, d_bc_* when presented with sequences (*a, b, c*), it should return *d_ac_, d_bc_, d_ab_* when given (*c, a, b*) as input. This is achieved by exploiting the fact that self-attention is naturally a permutation-equivariant function of a set of objects —permuting the elements in the input set yields the same permutation on the outputs. The fact that our network’s output, given a permutation of the sequences in the input MSA, corresponds simply to the induced permutation of the pairs, is then guaranteed by the choice of a symmetric representation (simple average) of pairs and by the fact that all the fully connected layers are position-wise, acting identically on each position in each pair (or sequence in the first layer). Phyloformer hence sees each permutation of an MSA as the same data point, which leads to the same predictions and parameter updates during training.

Furthermore our network is invariant to permutations of the sites in the MSA, which allows us to exploit the assumption that the process of evolution at each site is independent and identically distributed so that permuting the sites (the columns in the alignment) we simply get another realization of the same evolution process. Indeed the network structure guarantees in the same way equivariance under permutation of the columns until the last step, where taking the mean over these we ensure the output’s invariance. It is important to notice nevertheless that this doesn’t limit the applicability of the proposed network to scenarios in which a more sophisticated model of evolution, which models interactions between sites, is considered. In such a case it would suffice to add a positional encoding, as in classical transformers (Vaswani et al., 2017), to the network’s input without the need to modify its structure.

## 5 Results

### 5.1 Datasets

In this work we generate datasets using two different sequence simulators, Seq-Gen (Rambaut, 2017) and Evosimz (Zou et al., 2020).

The first model of evolution we tested our method on is a standard one, based on the PAM exchangeability matrix (Dayhoff et al., 1978), widely used in the phylogenetics community, and implemented in several phylogenetics tools (Rambaut, 2017; Huelsenbeck and Ronquist, 2001; Yang, 2007). We simulated trees as in Zou et al. (2020), *i.e.*, for each *n,* random *n*-leaves tree topologies were generated with the ete3 (Huerta-Cepas et al., 2016) populate() function and branch lengths were drawn uniformly between 0 and 1. The corresponding alignments were then obtained by simulating evolution along the branches with the PAM evolution model implemented in the software Seq-Gen (Rambaut, 2017). We used such simulations to train Phyloformer and investigate the effect of the number of leaves or the number of sites on the performances of various pieces of software.

The second model was presented in Zou et al. (2020) and implemented in their software Evosimz. It is based on a model that combines 9 different amino acid replacement matrices, a Gamma distribution for rate heterogeneity across sites, and site-specific amino acid equilibrium frequencies across sites. Further, the model can simulate heterogeneity across branches in both the rate of sequence evolution and equilibrium frequencies. In this simulator, the branch lengths found on the trees do not exactly correspond to expected numbers of per site substitutions, see Zou et al. (2020). However, this peculiarity has no consequence over our evaluation of the reconstruction methods, because we rely on Robinson-Foulds distances between true and estimated phylogenies that do not depend on branch lengths. We relied on existing datasets that make phylogenetic reconstruction challenging (Zaharias et al., 2022). These datasets have short sequences (200 amino acids) and long underlying branch lengths. Zaharias et al. (2022) looked at three different combinations of parameters (with distinct branch length distributions and amounts of time heterogeneity). For each of these parameter configurations they moreover considered different branch lengths distributions (corresponding to increasingly difficult phylogenetic estimation), multiplying by 10, 100 and 1000 their average, obtaining therefore 12 test datasets in total, each made of 20 trees along with the corresponding alignments containing 20 sequences of length 200. These datasets allowed us to evaluate the applicability of our method to a fairly complex model and to directly compare our results to those reported in Zaharias et al. (2022).

### 5.2 Performance on alignments simulated using Seq-Gen

#### 5.2.1 Reconstruction accuracy on alignments simulated using Seq-Gen

We trained our network on 100,000 alignments generated using Seq-Gen (Rambaut and Grass, 1997) (see section 5.1) with *n* = 20 and a sequence length of *L* = 200. We then generated additional simulations to investigate the effect of the number of leaves or the number of sites on phylogenetic reconstruction accuracy and on the resources required by various pieces of software. To investigate the effect of the number of leaves, we generated 1500 alignments of length 200 sites, varying the number of leaves between 10 and 100 by increments of 10. To investigate the effect of the number of sites, we generated datasets with 30 leaves, but varied sequence length between 200 and 1500 by increments of 100. In both cases, 150 tree-alignment pairs were generated for each parameter setting. We compared our method to the IQ-TREE (Minh et al., 2020) implementation of maximum likelihood, and to a standard distance method implemented in the R phangorn package (Schliep, 2011) which computes the maximum likelihood estimates of pairwise evolutionary distances and then reconstructs a tree using NJ (Saitou and Nei, 1987). Both methods were run with the correct model of sequence evolution. Figure 4 shows the normalized Robinson-Foulds distance (Robinson and Foulds, 1981) between the true and reconstructed trees for all three methods. Phyloformer performs better than the standard distance method across all parameter settings, and rivals with maximum likelihood in settings that most closely resemble its training data: when the number of leaves is close to 20, and when the number of sites close to 200. The degraded performance of Phyloformer on larger trees could be explained by the fact that larger trees have a longer average path length between leaves, which results in longer distances (given by the cumulative branch lengths along the path). These long distances can fall outside the range of distances seen by the network during training. The issue could thus be overcome by training Phyloformer on larger trees and/or with a wider range of branch length distributions. Similar results hold under mild model misspecification, namely in the situation where the data has been generated under the WAG model of sequence evolution, but maximum likelihood inference is still performed with PAM, and Phyloformer was trained on data generated using PAM (Sup. Figure 2).

**Figure 4:**
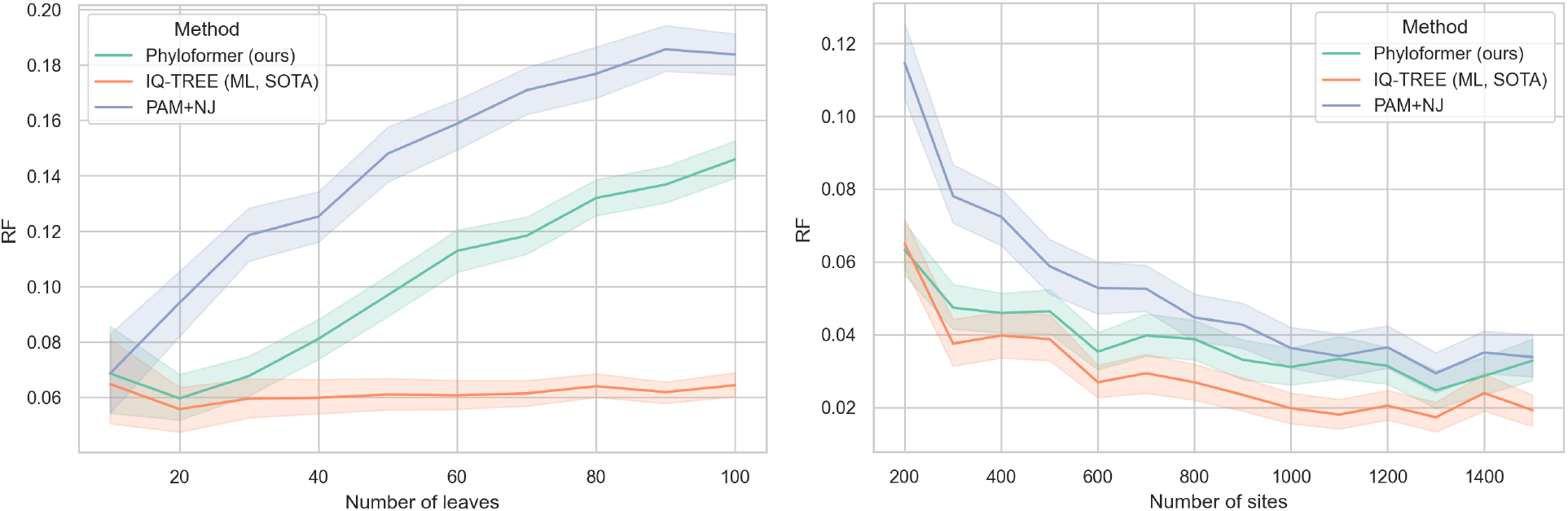
Mean performances of IQ-TREE (ML, State Of The Art, orange), PAM+NJ (distance method, blue), and Phyloformer (ours, green) on datasets varying in the number of leaves (left) or the number of sites (right). Shaded areas correspond to 95% confidence intervals of the mean. 150 trees were reconstructed for each number of leaves or each number of sites.

### 5.3 Performance on alignments simulated using Evosimz

We relied on alignments simulated using Evosimz (Zaharias et al., 2022) to assess the performance of several phylogenetic reconstruction methods on data simulated using a sophisticated model (see section 5.1).

To test the robustness of our model we trained Phyloformer only on alignments generated with the configuration resulting in the easiest simulations among the 12, i.e. the one in which all methods perform best (corresponding to the test dataset in the upper left corner of Figure 5). As above, we used a training dataset of 100,000 alignments with *n* = 20 and *L* = 200.

**Figure 5:**
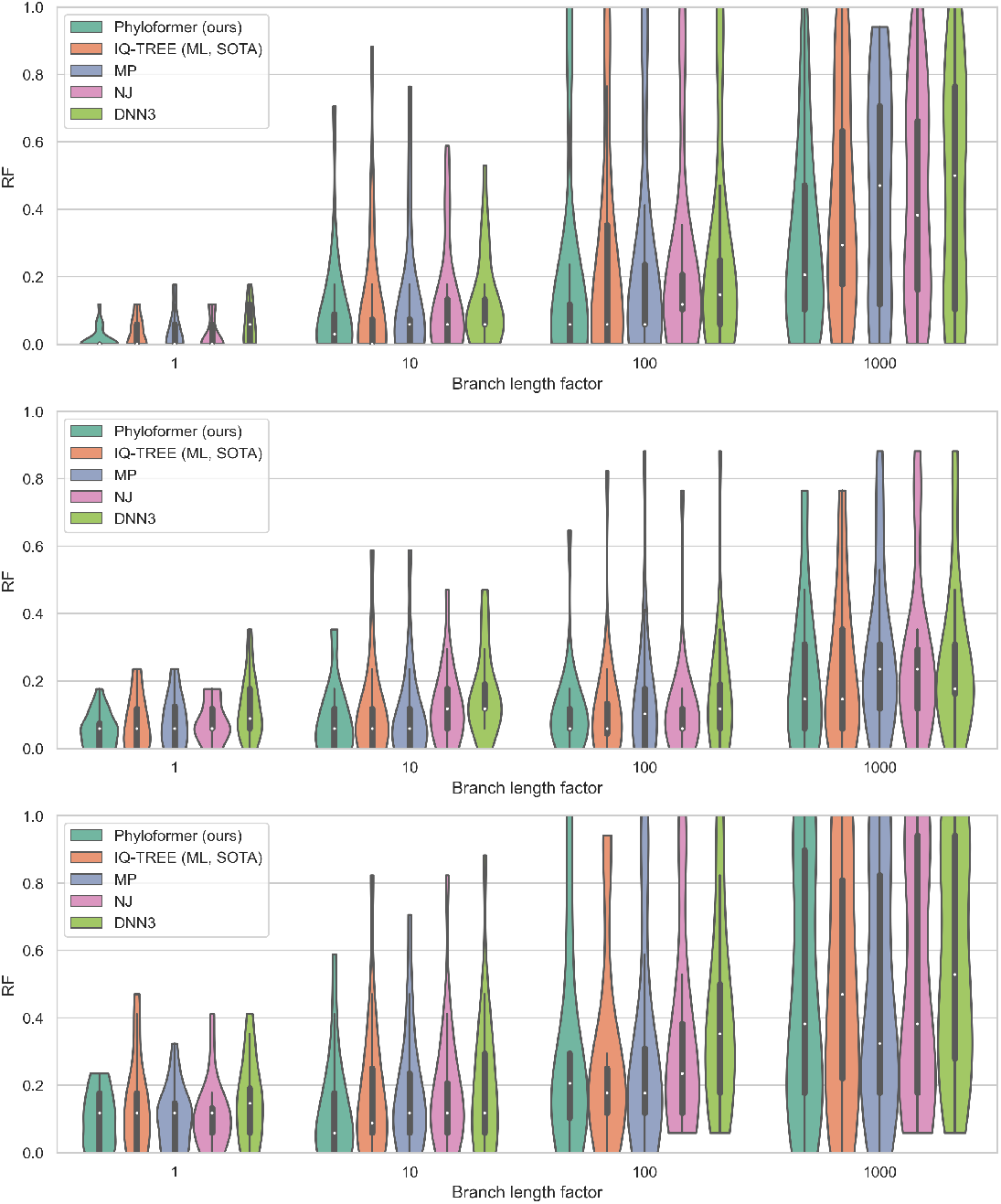
Violin plots of normalized Robinson-Foulds distances between the true trees and the trees estimated by 5 different methods, on datasets generated with different sets of parameters which reflect different difficulties in phylogenetic reconstruction. Results are shown for DNN3, IQ-TREE with ModelFinder, Maximum Parsimony, NJ (with hamming distances), and Phyloformer. Although our network was trained only on simulations generated with the easiest set of parameters (first dataset, upper left corner) it manages to always concentrate more mass towards low RF distance values than all other methods. As a result it outperforms them all in 9 out of 12 test datasets in terms of mean RF distance (Table 1).

In Figure 5 we show the violin plots of the Robinson-Foulds distances between real and predicted tree topologies, for all 12 datasets in Zaharias et al. (2022). We evaluated 5 different methods: Phyloformer, IQ-TREE with ModelFinder (which chooses the best reconstruction model according to the Akaike information criterion) (Nguyen et al., 2015), Maximum Parsimony (Farris, 1970; Fitch, 1971), Neighbor Joining (with simple hamming distances), and Zou et al. (2020)’s neural network DNN3. To avoid cluttering the plot we omitted the other methods analyzed in Zaharias et al. (2022), namely different implementations of maximum likelihood (performing worse overall than the one here reported) and the other two Zou et al. (2020)’s networks. In Zou et al. (2020), DNN3 performed better overall and was published as a ready-to-use phylogenetic inference tool. Nevertheless, a table reporting the mean normalized RF distance for all the methods can be found in the Supplementary Material. All methods perform best on the easiest simulations, on the left side of the plots. As reported in Zaharias et al. (2022), the neural network DNN3 does not perform as well as the other reconstruction methods on these simulations. Maximum Parsimony and the distance method often perform better, but the best performing methods are the maximum likelihood approach IQ-TREE, and our method Phyloformer. For Phyloformer the distribution is systematically more skewed towards low RF distance values than for IQ-TREE. Further, if we only look at the mean RF distance our method performs better than all other methods on 9 out of the 12 datasets (Table 1). These results show that Phyloformer can perform better than Maximum Likelihood when the data has been simulated under complex models of sequence evolution.

#### 5.3.1 Speed and memory requirements on alignments simulated using Seq-Gen

For IQ-TREE (SOTA), Phyloformer and PAM+NJ we evaluated the memory and time taken to analyze all 150 trees for a given number of sites and number of sequences. All methods were limited to using a single CPU core, and Phyloformer was allowed to use the GPU. Time and memory footprints were measured in triplicates on the Seq-Gen datasets, using the benchmark options in snakemake (Mölder et al., 2021) on a machine with an Intel Core i7-10700 processor with 32GB of RAM and equipped with an Nvidia RTX A4000 GPU. Memory footprint for Phyloformer, which uses the GPU, were obtained with a call to `torch.cuda.max_memory_allocated` in pytorch (Paszke et al., 2019). All methods increase their footprint as the number of sequences or sites grow. Phyloformer has the largest memory usage (Sup. Figure 2), and requires almost 2.9GB for alignments with 1,500 sites and 30 sequences while IQ-TREE, which has the lowest memory usage, only requires 16.5MB. In terms of speed however (Figure 6), IQ-TREE is the slowest, requiring up to ~5000s for the 150 alignments with 100 sequences, or up to ~7500s for the alignments with 1,500 sites. PAM+NJ is the fastest, and requires ~26s and ~5.7s, respectively, while Phyloformer requires ~ 174s and ~110s. Phyloformer is thus up to 68 times faster than IQ-TREE in our hands, when run on alignments with 30 sequences and 1,500 sites. Furthermore it is worth noting that here each alignment was processed as a single batch of size one by our network while the method lends itself to a high degree of parallelization being able to process multiple alignments in the same batch, which can significantly improve its speed performances, the only constraint being the available memory.

### 6 Discussion and future work

In this manuscript we have presented Phyloformer, a new method for phylogenetic reconstruction using deep learning. We have showed on two types of simulations that it could approach or beat state-of-the-art (SOTA) maximum likelihood methods that use all the information available in the alignment (Nguyen et al., 2015) while being significantly faster. Our use of two simulators reduces the risk of a simulator-related bias in our conclusions. The simulations performed using the simplest model revealed two regimes: on alignments with numbers of sequences similar to the one used to train the network (20) Phyloformer lead to Robinson-Fould distances that were very close to the ones obtained by IQ-TREE while being one order of magnitude faster. For example, on alignments of 30 sequences of length 200 produced with Seq-Gen (right panel, Figure 6), Phyloformer obtained a mean RF distance of 0.063 in 9s, whereas IQ-TREE took 152s to obtain a mean distance of 0.065. Performances worsened progressively as we drifted away from the setting used to generate training data by increasing the number of leaves, but always remained better than the performance of model-based estimates of pairwise distances between sequences combined with the NJ clustering algorithm (PAM+NJ). In this less favorable regime, Phyloformer was up to 25 times faster than IQ-TREE even for sequences of length 200, and 7 times slower than PAM+NJ, therefore offering a trade-off between speed and accuracy. We are confident that this less favorable regime can be alleviated by using more diverse settings for training, including trees with more leaves.

**Figure 6:**
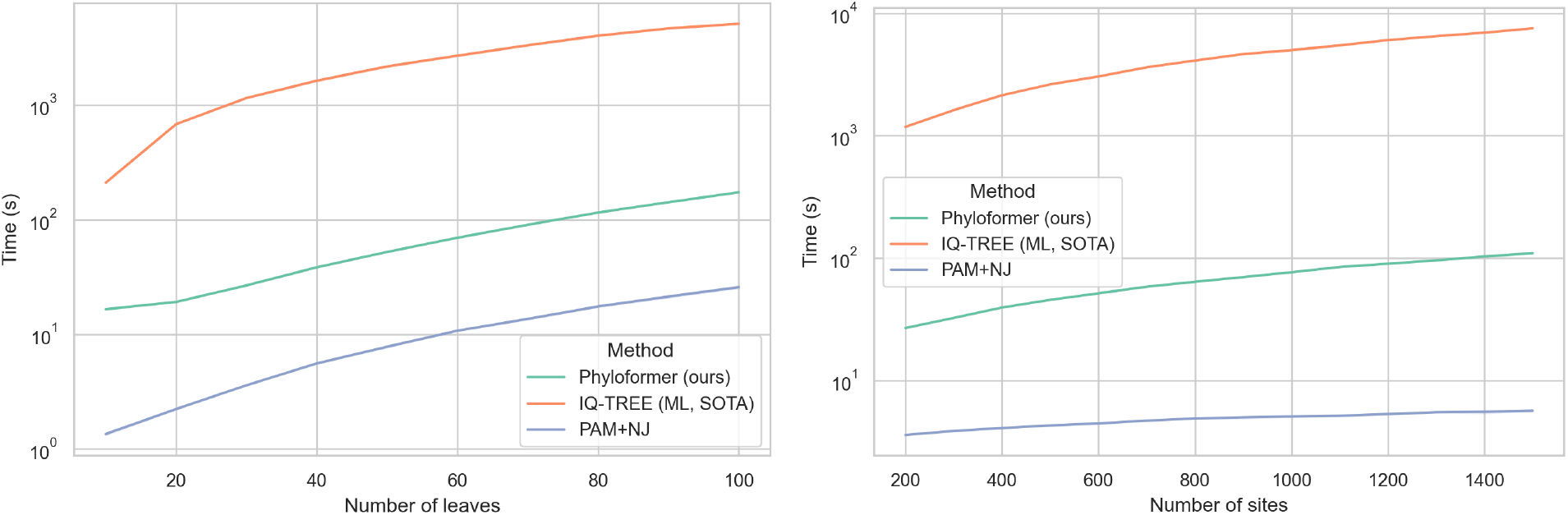
Time footprints, on a logarithmic scale, of IQ-TREE (ML, State Of The Art), PAM+NJ, and Phyloformer on datasets of 150 trees varying in the number of leaves (left) or the number of sites (right). Colors as in Figure 4.

On the simulations obtained with a more complex model implemented in Evosimz, where the number of sequences and sites remained the same between training and testing, Phyloformer performed as well or better than all methods, IQ-TREE included. In particular, Phyloformer outperformed existing deep learning approaches for phylogenetic reconstruction. These results show that Phyloformer can perform as well as a state-of-the-art maximum likelihood method on complex simulations. Although Phyloformer was trained on data generated using the same Evosimz simulator, the training datasets were all generated using the easiest parameter settings that we have tried. This, together with the results obtained on WAG simulations (Sup. Figure 2), indicates that Phyloformer has some robustness against model misspecification. Its robustness to more radical misspecifications (e.g., branch length or topology distributions not seen during training) will require additional analyses. Similarly, several processes affecting empirical data have been kept out of the current simulations. For instance, GC-biased gene conversion Ratnakumar et al. (2010) or mutational biases (e.g., CpG hypermutability, Meunier et al. (2005)) have not been included. It is unclear how the various methods would fare on data simulated with models that reproduce these processes. Overall, precisely evaluating the level of robustness of Phyloformer will still require some work, and will be a necessary step to making it usable as a routine tool. Depending on the possibility of a single network to provide robust predictions in all settings, one may need to rely on several ones or to offer the possibility to fine-tune to a specific situation. Importantly, future versions should also propose a measure of uncertainty along with the predictions (Gal and Ghahramani, 2016; Charpentier et al., 2022).

A natural follow-up question is whether Phyloformer could perform as well as maximum likelihood methods on empirical datasets. This question is difficult to answer with empirical data, because the true phylogeny is rarely known without a doubt. This is particularly true for gene trees, which can differ from the species tree due to gene duplication, loss, transfer, or incomplete lineage sorting Szöllõsi et al. (2015). However, in cases where the species phylogeny is known, one could count the proportion of reconstructed gene trees or subtrees that match the species phylogeny. Another way to answer this question would be to use realistic simulations, where the generating phylogeny is known, and where the model of sequence evolution has been shown to generate datasets that are realistic, *i.e.*, very hard to distinguish from empirical datasets. In future work, we will use generative models (Kingma and Welling, 2014; Goodfellow et al., 2014; Yelmen et al., 2021), relying on an architecture similar to Phyloformer, to assess how realistic current simulations are and to produce more realistic ones. A more realistic simulator of sequence evolution will allow for more accurate evaluations of reconstruction methods and conceivably lead to better performances on real data of our network, retrained on such more true to life simulations.

Phyloformer relies on two technical innovations to learn how to accurately reconstruct phylogenies from MSAs. Firstly, it adresses phylogenetic reconstruction as a regression problem. Previous approaches relied on a classification task between a number of competing tree topologies. The number of tree topologies grows super-exponentially with the number of leaves, which makes such an approach unable to scale to MSAs with more than a few sequences.

Secondly, through a transformer-based architecture, Phyloformer uses self-attention, both across sites and pairs in the MSA. Self-attention provides an expressive mechanism that models how each pair (resp. site) should inform the others within the MSA. This mechanism is compositional: it relies on learnable functions that each apply to individual objects (pairs or sites) and whose outputs are assembled to determine how information is shared. This compositional structure also makes the resulting function permutation-equivariant and able to process MSAs of varying sizes: Phyloformer learns how to predict the evolutionary distance between a pair of sequences given all other sequences in the MSA.

The combination of these two technical innovations—the regression formulation and self-attention mechanism— produces an effective mean to jointly predict all pairwise distances across the MSA. At each stage of the learning, every pair of sequences benefits from the most up-to-date information extracted from all others, not just their raw sequences. As previously pointed out (Zaharias et al., 2022), using global information is key to an accurate reconstruction. Future efforts will include the exploration of more sophisticated invariant functions rather than the simple average both for the pair representation and the final column pooling as well as an analysis of the performances of the network combined with different distance-based methods other than simple NJ, such as BioNJ.

More generally, the performance of Phyloformer could be improved by simulating from arbitrarily complex models of sequence evolution, with slight changes to the structure of the network. We expect this to increase its edge against likelihood-based methods, whose computational cost can become intractable unless simplifying assumptions are made. In particular, the current Phyloformer model is invariant by permutation of the sites to take advantage of the site independence in the simulation models, but could easily be modified to account for heterogeneity of the evolutionary process along the sequence or to consider site dependencies, using positional information—a standard approach in the transformers literature. Similarly, accounting for insertion-deletion processes is known to be a hurdle for likelihood methods (Redelings and Suchard, 2007), and would be straightforward using our paradigm. Phyloformer could even be extended to unaligned sequences by discarding the site dimension with global pooling earlier on in the network. A learning approach that bypasses the alignment step and yet yields accurate phylogenies could be extremely useful, as aligning sequences is computationally costly.

Currently Phyloformer applies self-attention layers to pairs of sequences, making its scaling quadratic in the number of tree leaves. A clear improvement would be to use self-attention on single sequences and use the final embeddings to estimate pairwise distances—in the spirit of MSA Transformer (Rao et al., 2021). Our initial attempts in this direction have led to poorer accuracies when reconstructing phylogenies over more leaves than what was seen in the training data. Some more effort on the design of the network and training data could solve this issue and allow for a more scalable version of Phyloformer, requiring less memory. Our method will also benefit from the growing literature on faster self-attention and more efficient versions of transformers (Tay et al., 2020; Dao et al., 2022).

We have demonstrated for the first time that a neural network can achieve state-of-the-art performance for phylogenetic reconstruction, in a significantly reduced amount of time. This was observed when test data and training data share similar numbers of leaves, under commonly used models of sequence evolution. In the future, we will improve our training data to extend the applicability of Phyloformer. With training data that covers enough of the variation observed in empirical data, we expect that Phyloformer will offer performance rivaling ML across the board. The ability to perform accurate phylogenetic inference at a much lower computational footprint could change practices in molecular evolution, comparative genomics, epidemiology, and metagenomics. Beyond phylogenetic reconstruction, these results suggest that neural networks will be able to compete with ML and Bayesian approaches for reconstructing dated phylogenies, inferring gene trees and species trees, and identifying sites and genes under selective pressure. It is quite possible that, ten years from now, most practitioners will be using faster and better methods, based on neural networks, that are currently under development.

## Code and data availability

The code for Phyloformer, the pretrained models, and all the datasets analyzed in this work can be found at https://github.com/lucanest/Phyloformer.

## Acknowledgements

The authors thank Dexiong Chen, Martin Ruffel, Johanna Trost and Philippe Veber for insightful discussions.

This work was funded by the Agence Nationale de la Recherche (ANR-20-CE45-0017). It was granted access to the HPC/AI resources of IDRIS under the allocation AD011011137R1 made by GENCI. The taxon silhouettes in Figure 2 are modified from public domain images in the PhyloPic database.

## Footnotes

2 The position-wise fully connected layers in the architecture act likewise, one can interpret them as simple 1 × 1 2D convolutions.

