## Supplementary materials for "Phyloformer: towards fast and accurate phylogeny estimation with self-attention networks"

**Luca Nesterenko, Bastien Boussau,* Laurent Jacob***
Biometry and Evolutionary Biology laboratory (LBBE)
University Claude Bernard Lyon 1
Lyon, France


#### Contents

**1 Evosimz simulations results** **1**

**2 Model misspecification** **2**

**3 Memory usage** **2**

#### 1 Evosimz simulations results

| method<br>experiment | dnn1 | dnn2 | dnn3 | ghost | iqtree | mp | nj | phyloformer | upgma | wag |
|---|---|---|---|---|---|---|---|---|---|---|
| configuration1_1 | 0.059 | 0.071 | 0.059 | 0.050 | 0.035 | 0.038 | 0.029 | **0.015** | 0.162 | 0.032 |
| configuration1_10 | 0.126 | 0.162 | 0.126 | 0.135 | 0.109 | 0.118 | 0.121 | **0.091** | 0.271 | 0.141 |
| configuration1_100 | 0.276 | 0.285 | 0.259 | 0.221 | 0.226 | 0.225 | 0.241 | **0.185** | 0.350 | 0.212 |
| configuration1_1000 | 0.450 | 0.421 | 0.450 | 0.435 | 0.409 | 0.441 | 0.426 | **0.297** | 0.509 | 0.479 |
| configuration2_1 | 0.132 | 0.097 | 0.121 | 0.094 | 0.079 | 0.084 | 0.079 | **0.056** | 0.188 | 0.085 |
| configuration2_10 | 0.179 | 0.138 | 0.165 | 0.121 | 0.109 | 0.099 | 0.126 | 0.088 | 0.312 | **0.076** |
| configuration2_100 | 0.168 | 0.147 | 0.168 | 0.129 | 0.118 | 0.146 | 0.118 | 0.112 | 0.265 | **0.106** |
| configuration2_1000 | 0.321 | 0.288 | 0.265 | 0.279 | 0.238 | 0.282 | 0.268 | **0.221** | 0.412 | 0.297 |
| configuration3_1 | 0.165 | 0.135 | 0.150 | 0.129 | 0.138 | 0.104 | 0.112 | **0.100** | 0.279 | 0.135 |
| configuration3_10 | 0.200 | 0.188 | 0.200 | 0.182 | 0.171 | 0.174 | 0.182 | **0.126** | 0.291 | 0.188 |
| configuration3_100 | 0.371 | 0.362 | 0.400 | 0.303 | 0.291 | 0.294 | 0.326 | **0.279** | 0.512 | 0.329 |
| configuration3_1000 | 0.556 | 0.556 | 0.568 | **0.474** | 0.485 | 0.482 | 0.506 | 0.479 | 0.650 | **0.474** |

Table 1: Mean normalized RF values for different reconstruction methods, evaluated on the 20 trees datasets generated with the Evosimz simulator (Zou et al., 2020) by Zaharias et al. (2022). In bold the minimum mean reconstruction error among all the methods for each dataset.

In the table we compare the performances, in terms of mean normalized RF distance of all methods considered in (Zaharias et al., 2022) against those of Phyloformer. We have maximum parsimony (mp), the three neural networks by (Zou et al., 2020), Neighbor Joining with Hamming distances (nj), UPGMA, and three different IQ-TREE implementations of maximum likelihood, using ModelFinder (iqtree) and using the models GHOST and WAG. Our method attains the lowest mean error in 9 out 12 parameter settings being only slightly outperformed by slower maximum likelihood methods in the remaining 3.

*Equal contribution

### 2 Model misspecification

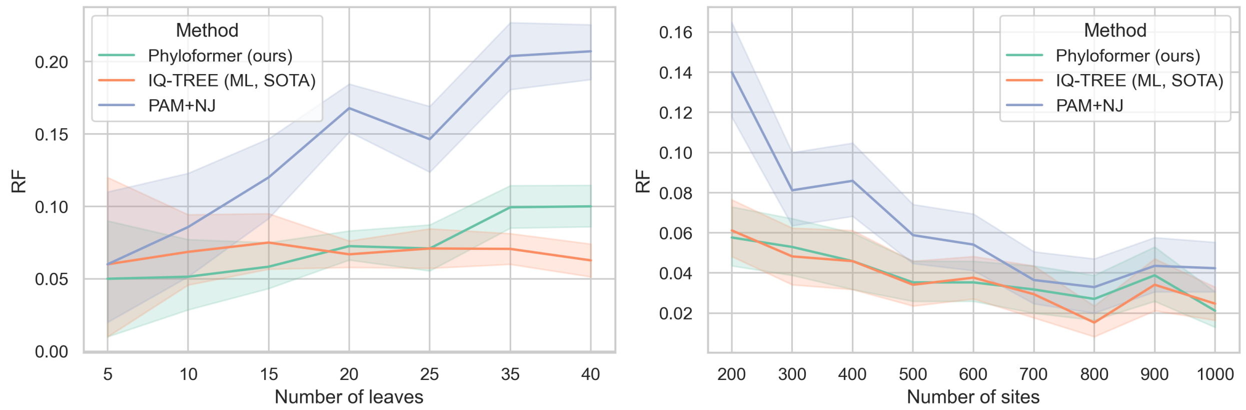


Figure 1: Normalized RF distances for different methods under model misspecification.

To test to some extent the robustness to model misspecification of Phyloformer we compared against maximum likelihood and a standard distance method providing to all three an incorrect model of evolution. Namely we simulated alignments with the software Seq-Gen under the same scheme as in 5.2.1, but under the WAG model of amino acid substitution. We then compared the performances of our network (trained on alignments generated under the PAM model, *i.e.*, the same as in Section 5.2.1) against those of maximum likelihood and of a standard distance method, both using PAM. 50 alignments were generated for each configuration of $n$ and $L$, keeping $L = 200$ fixed while increasing the number of sequences $n$ from 5 to 40 and keeping $n = 20$ fixed while increasing the number of sites $n$ from 200 to 1000. We can observe in Figure 2 the same trend as with the PAM simulations, with Phyloformer consistently outperforming the standard distance method and having accuracy comparable to that of maximum likelihood, with degrading performances as the value $n$ gets progressively bigger than the one the network has been trained with.

### 3 Memory usage

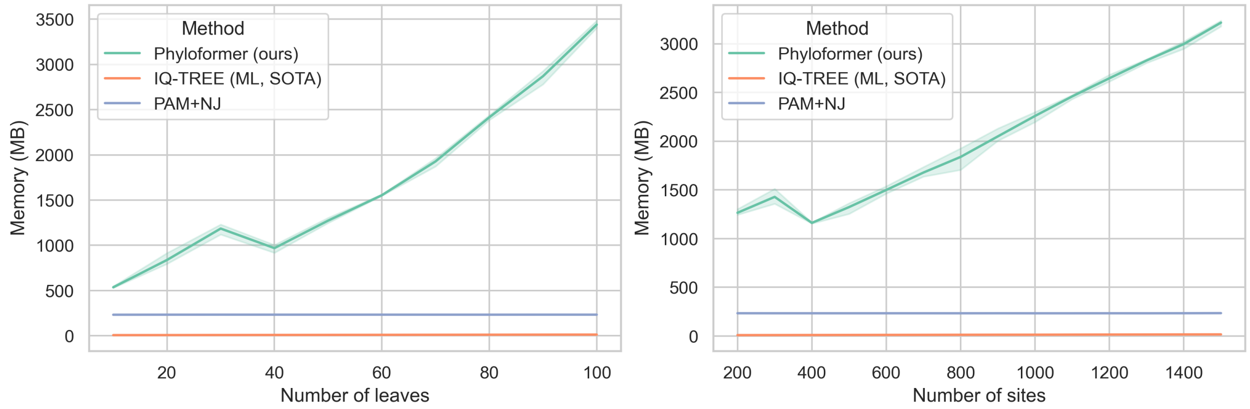


Figure 2: Memory usage of IQ-TREE (ML, State Of The Art), PAM+NJ, and Phyloformer on datasets varying in the number of leaves (left) or the number of sites (right). Colors as in Fig. 4

.

The memory usage of Phyloformer as shown Figure 2 (see also section 5.3.1) is linear in terms of the number of sites, but quadratic in terms of the number of leaves. This remains a drawback of our method, limiting the applicability to very large trees and the possibility of processing several trees in parallel with larger batches.
